# Long-range linkage maintained male-specific loci on a young Y chromosome prior to recombination shutdown

**DOI:** 10.64898/2026.08.31.748364

**Authors:** May Wang, Mohammadebrahim Akhavizadegan, Jen-Yu Wang, Ching-Ho Chang, Kevin H-C Wei

## Abstract

Suppression of recombination on the Y chromosome maintains linkage between male-specific loci and is commonly established by inversions. Here, we describe a young, inversion-free Y chromosome (neo-Y) in *Drosophila albomicans* with a unique history and paradoxical signature of exchange. Prior to recombination shutdown in males, it repeatedly recombined with the X-linked counterpart (neo-X) but at the same time preserved complete long-range linkage of the chromosome ends. By assembling multiple neo-Ys chromosomes and QTL-mapping, we show that double crossovers maintained linkage between the male-sex determining region at one end to sexually antagonistic alleles and a locus essential for spermatogenesis at the other. We argue that such long-range linkage of distal sex-specific loci disfavors inversions but instead encourages the emergence of achiasmy.

## Introduction

Cessation of recombination is critical to initiate the divergence of sex chromosomes. Transmitted strictly through males, the Y chromosome is expected to carry both the male-determining and male-beneficial loci (MBL). If the Y continues to recombine with the X, unfit sons lacking the MBL will be sired and if the loci are further sexually antagonistic (SA), unfit daughters with female-detrimental alleles will also be produced, further enforcing the deleterious impact (*1*). Natural selection then favors the cessation of recombination locking the linkage of the male determining locus to MBL on the Y. Once exchange stops, the Y chromosome will progressively degenerate due to a combination of evolutionary forces including hitchhiking and Muller’s ratchet, resulting ultimately in a gene-poor and repeat-rich chromosome [reviewed in (*2, 3*)].

Linkage of the male determining locus to MBL typically proceeds in a stepwise fashion that progresses across the chromosome over evolutionary time [reviewed in (*2*–*4*)]. The male-determining locus and nearby MBL become locked by local suppression of recombination while the remainder is free to recombine with the X. Suppression can be achieved by a single chromosomal inversion although other mechanisms are possible (*5*). To capture additional MBL or SA loci that arise, successive inversions can expand the nonrecombining region, producing less degenerate segments with more recent history of exchange (*5, 6*). These regions are known as evolutionary strata and observed on Y chromosomes in mammals (*7, 8*), fish (*9, 10*), insects (*11, 12*), and plants (*13, 14*).

*Drosophila* has long been a key genus for understanding sex chromosome evolution due to multiple independent emergence of neo-sex chromosomes. Because males are, typically, achiasmate, suppression of recombination is thought to be immediate once a chromosome becomes male-specific, i.e. a neo-Y chromosome. Not unique to flies, sex-specific achiasmy has independently evolved over 30 times (*15*) and always in the heterogametic sex (e.g. XY males and ZW females), adhering strictly to the Haldane-Huxley rule (*16, 17*). It has been proposed that achiasmy emerged to prevent recombination between sex chromosomes in the heterogametic sex with the autosomes as a pleiotropic collateral. However, because crossovers are often required for the fidelity of chromosome segregation, genomewide recombination shutdown presents an evolutionary paradox. The deleterious potential of achiasmate meiosis is particularly salient in *Drosophila*, which has evolved chromosome-specific pairing systems to ensure proper disjunction (*18*–*21*) that were either prerequisite for achiasmy to spread or adaptation that followed. Given that inversions are commonplace and can create nonrecombining stratum without compromising segregation fidelity, why selection would favor global recombination shutdown via achiasmy remains unclear .

Here, we investigate a *Drosophila* species harbouring a young neo-Y chromosome with a history of frequent exchange with its neo-X counterpart – a scenario made possible because the species originated from a lineage with males recombination (*22*). We discover that despite collinearity with the neo-X and repeated COs along the arm, the distal ends of the neo-Y remain in tight linkage generating an unusual pattern of long-range linkage disequilibrium (LD). We show that this signature resulted from double COs that maintain linkage of the male-determining locus at one end with MBL and SA loci at the other. Because the species subsequently reverted to male achiasmy, we propose that maintenance of long-range LD may be a condition that favors the global shutdown of recombination as opposed to forming nonrecombining strata via inversions.

### Long-range LD on the neo-Y despite repeated historical recombination

The neo-sex chromosomes of *D. albomicans* originated less than 200 kya (*23*) from Robertsonian fusions between the autosome Chr 3 (a.k.a. Muller CD) to the ancestral X (a.k.a. Muller A) and Y (Fig. 1A) (*24*). Despite a single origin of the neo-Y (Fig. S1A) (*25*), it has repeatedly recombined, via historical male recombination with the neo-X (*22*), creating several different haplotypes distal to the centromere. Previously, three geographically distributed haplotypes (Y1, Y2, and Y3) were identified (*26*), but there appears to be additional exchange even within each haplotype group (Figure 1B). In fact, there are up to six neo-Y haplotypes towards the middle of the chromosome (Figure 1C), suggesting recombination with the neo-X was commonplace. However, the capacity for male recombination appears to have ceased as no recombination occurs in male test crosses (*22*) or F1 hybrid males (see cross scheme below) and there is no evidence of on-going exchange in inbred strains which should homogenize genotypes between the neo-sex chromosomes (fig. S2).

**Figure 1.**
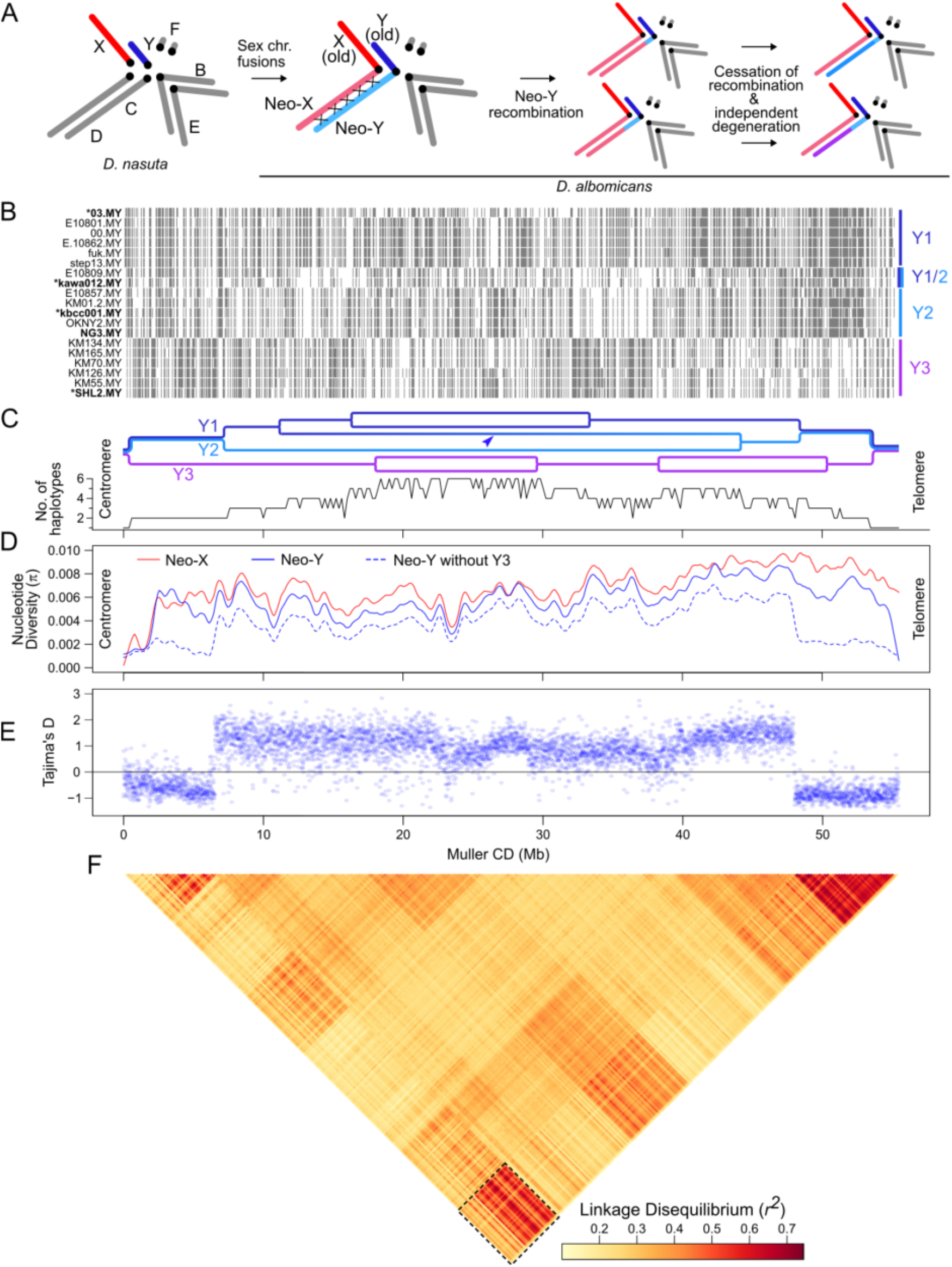
Long range linkage of the ends of the neo-Y despite repeated recombination in *D. albomicans*. **A**. Evolution history of the neo-sex chromosomes in *D. albomicans*. **B**. Genotype and haplotype representation across different neo-Ys. White and gray bars represent reference and nonreference SNPs, respectively. Genotypes are subsetted such that 50 positions with SNPs are randomly selected every 10kb. Positions based on homologous positions on the neo-X. **C**. Top, schematic representation of haplotype structure across the neo-Y. Lineage bifurcations represent positions of COs that caused haplotype switches and introduced novel haplotypes from the neo-X onto the neo-Y; lineage convergences represent COs that eliminated haplotypes. Arrowhead points to the neo-Y haplotype that is a mixture of both Y1 and Y2. Bottom, number of haplotypes in each 100kb window are inferred based on the number of monophyletic clades the neo-Ys form on the phylogeny; for representative phylogenetic trees, see fig. S1. **D-E**. Chromosome-wide nucleotide diversity (**D**) and Tajima’s D (**E**) of the neo-X and neo-Y chromosomes. **F**. Chromosome-wide linkage disequilibrium inferred from neo-Y genotypes; *r*^*2*^ averaged across SNPs within 10kb windows. Signature of long-range LD in dotted box.

Because of the frequent exchange through ancestral male recombination, neo-Y’s nucleotide closely tracks neo-X’s and remains highly elevated outside of the centromere proximal region (Fig. 1D and E) which has low diversity due to the strong bottleneck imposed by the single fusion (fig. S1A) and centromeric suppression of recombination (*27*). Accompanied by sharp rises in nucleotide diversity, CO at ∼2Mb distinguishes Y3 from Y1 and Y2 and another at ∼5Mb separates Y1 and Y2 (Fig. 1B and C); additional exchanges occurred throughout the middle of the chromosome arm, including some that created a chimeric Y that, curiously, is a mix of Y1 and Y2. Notably, Y3 further exhibits evidence of introgression from a closely related species near the distal end (fig. S1C). While diverse haplotypes have been introduced through COs, the distal end of the neo-Y unexpectedly displays signatures of the same bottleneck as the centromere-proximal end, showing depleted diversity and monophyly consistent with a single origin (Fig. 1C,D and fig. S1F). This is particularly noticeable when only Y1 and Y2 genotypes are considered, as the two lack introgression like Y3 that would introduce excess diversity.Strikingly, the two ends of the chromosome arm are in high linkage disequilibrium (LD) (Fig. 1E), which is also reflected by the collapse of haplotype diversity towards the distal end of the chromosome back down to one (Fig. 1C). Therefore, even with repeated exchange across the middle, the two ends of the neo-Y remain, paradoxically, fully linked.

### No structural variation on the neo-Y to suppress recombination

To determine the source of the long-range LD, we first explored the possibility that structural rearrangements on the neo-X or neo-Y created physical proximity of the genetically linked ends. We used Nanopore long read sequencing to generate complete, phased assemblies (*28*) of four neo-Y chromosome arms (Fig. 2A, fig S3, and Supp table 1). Synteny analysis of gene order revealed that the neo-Ys are completely collinear with the neo-X, ruling out the possibility of suppression of recombination via inversions or rearrangements even though they are common in this species group (Fig. 2A) (*25, 29, 30*). Additionally, repeat content appears only moderately increased on the neo-Ys (Fig. 2B), ruling out the possibility of heterochromatin-induced suppression of recombination which is more likely for older Ys that accumulate more repeats (*5, 31*). Indeed, the neo-Y appears to show little signs of degeneration, with few gene losses and down-regulation (Fig. 2C, Fig. S3), the bulk of which are enriched at the ends where the neo-Y is most differentiated due to the lack of historical exchange (Fig. 2D). The absence of clear indicators of degeneration is unsurprising given the recent origin (*23*) and the repeated exchange with the neo-X in the middle which would have reverted any progress of degeneration by introducing freely recombining neo-X haplotypes yet to experience the degenerating forces (*32*). We conclude that long-range LD was maintained while the two chromosomes were unimpeded by rearrangements and fully capable of recombining.

**Figure 2.**
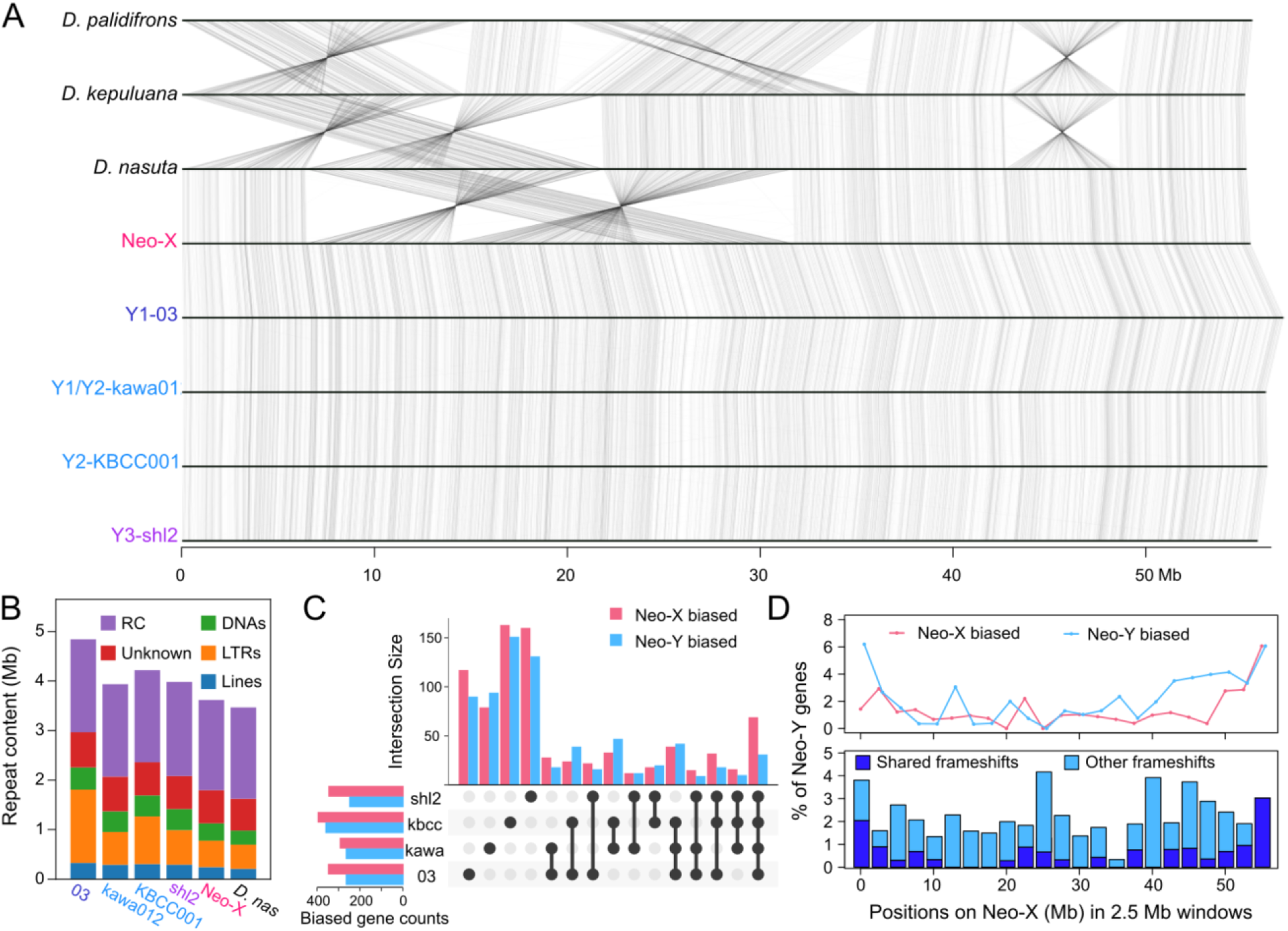
Chromosome-level assemblies of neo-Y haplotypes show collinearity and little degeneration. **A**. Synteny analysis of the neo-sex chromosomes and the ancestral autosome across *nasuta* subgroup species. Gray lines connect orthologous genes between chromosome assemblies. **B**. Repeat content on the neo-sex chromosomes. **C**. Accounting of haplotype-specific and shared genes with significantly differentially expressed neo-X (pink) and neo-Y (blue) gametologs in testes RNA-seq from males with different neo-Ys. **D**. Top, number of genes with significantly differentially expressed gametologs shared between males with different neo-Ys across the chromosome in 2.5Mb windows. Bottom, number of neo-Y gametologs with frameshifts shared between neo-Y assemblies across the chromosome in 2.5Mb windows. Positions are relative to the neo-X.

### Double crossovers maintain long range LD to locus essential for spermatogenesis

Single, or odd number, COs between collinear homologous chromosomes necessarily lead to haplotype switching of all distal loci and therefore cannot maintain linkage of distal ends. Given that every introduction of a new haplotype on the neo-Y collapses back to one (Fig. 1C), we consider the possibility that the observed long-range LD may have resulted from the persistence of double, or even number, CO recombinants, and hypothesize that the distal end of the neo-Y harbours a MBL. Double COs then preserve the linkage of the male determining locus on the old Y with the MBL on the neo-Y, while single CO recombinants are purged by selection due to the absence of the MBL.

To test this hypothesis, we took advantage of interspecies crosses with the sister species *D. nasuta* lacking the neo-sex chromosomes to generate a hybrid aneuploid female (*33, 34*) carrying the neo-X, neo-Y, and an unfused ancestral X (henceforth XXY females; Fig. 3A). Transmitted through female chiasmate meiosis, the neo-sex chromosomes can recombine with each other producing recombinant, matrilineally inherited neo-Ys (fig. S4C). The offspring were then genotyped by molecular markers at proximal, central, and distal locations on the neo-sex chromosomes. While we recovered no recombinant males with homozygous neo-Y genotypes indicating that several recessive lethal alleles have accumulated, males with homozygous neo-X genotypes were viable (Fig. 3B). Affirming our hypothesis, 1 CO recombinants were mostly sterile, significantly more compared to non-recombinants (p < 1e-10, Fisher’s exact test). Definitively, 2 CO males that maintained linkage of the distal ends have significantly higher fertility than 1 CO males (Fig. 3B; p = 0.000196, Fisher’s exact test). Therefore, a MBL essential for male fertility resides near the end of the neo-Y. Whole genome sequencing and QTL analysis of a subset of the recombinant males narrowed down the MBL down to a ∼3Mb region at the distal tip (Fig. 3C), although there are likely additional loci including hybrid incompatibilities (*35*) contributing to fecundity. Recombinant males lacking the MBL show clear defects in spermatogenesis compared to wildtype (Fig. 3D and E). Near the apical tip of the long testes, bundled sperm nuclei appear disordered (Fig. 3E’), elongate prematurely (Fig. 3E’’), and dissociate leading to individualized needle-like threads (Fig. 3E’, arrows) that are typically found near the seminal vesicle at the end of the testes track (*36*). These malformations likely lead to aborted spermatogenesis as no sperm can be found in the seminal vesicle (Fig. 3E’’’).

**Figure 3.**
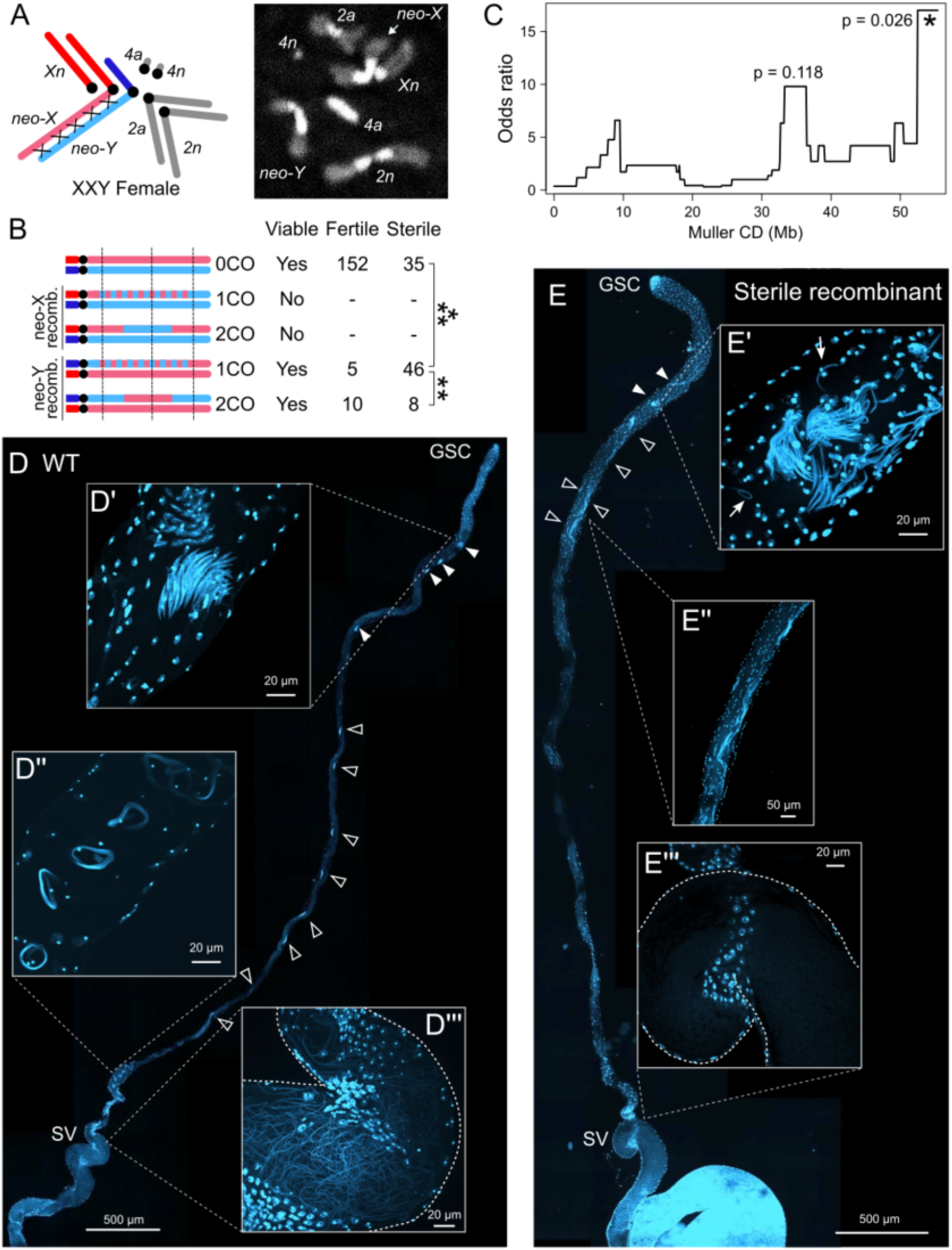
Recombinant mapping of male essential locus on the Neo-Y. **A**. Karyotype schematic (left) and mitotic chromosome spread (right) of XXY females. The *a* and *n* subscripts on chromosomes denote *D. albomicans* and *nasuta* chromosomes, respectively. **B**. Sons of XXY females crossed to *D. albomicans* males are PCR-genotyped at proximal, central, and distal (dotted vertical lines) regions using indel polymorphisms. Chromosome schematics indicate the inferred recombinant genotype based on the molecular markers. The fertility and viability of the males are tallied to the right. The nonrecombinant class includes males that inherited the Y from the mother and father. For all possible genotypic outcomes of the cross, see fig. S5. *** indicate significant difference between the 1CO and 0CO classes (p < 0.00001 Fisher’s Exact Test). ** indicates significant difference between the 2CO and 1CO classes (p < 0.0001). **C**. QTL association in odds ratio between neo-Y genotype across the chromosome and fertility of males with recombinant neo-Ys; see fig. S6 for genotypes of all sequenced individuals. D and E. WT (D) and sterile (E) testes stained with DAPI. GSC - germline stem cells; SV - seminal vesicles. Filled and blank arrow heads indicate sperm bundles before and during elongation, respectively. D’ & E’, Leaf and canoe stage spermatids; arrows point to individualized nuclei. D’’ - Coiling of individualized sperm bundles. E’’ Elongated sperm nuclei. D’’’ & E’’’ interior view of seminal vesicles.

### Sweeps of demasculinized alleles on the neo-X suggest resolution to sexual antagonism

Curiously, fecundity of the neo-Y-carrying XXY females decreased by more than 10-fold (Fig. 4A), a significant reduction (fig. S4C; p = 1.4e-11, Wilcoxon’s Rank Sum Test) even after accounting for the lethal aneuploid genotypes. Reduced fecundity is further accompanied by significant female-biased offspring sex ratio (Fig. 4B; p = 8.9e-13, Wilcoxon’s Rank Sum Test) likely due to both exposed recessive lethality on the neo-Y, and paucity of sons with matrilineal neo-Y possibly due to segregation distortion, but neither are sufficient to account for the drastic fecundity decline (fig. S7). As XXY aneuploidy is not intrinsically detrimental in *Drosophila* (*37*), our results suggest the neo-Y is deleterious to females. In the recombinant daughters of the XXY females, we further found associations between presence of the neo-Y genotype and reduced fecundity at the proximal and distal markers (Fig 4A). As all three regions show intermediate fecundity compared to the XX and XXY genotypes, there are likely multiple dominant SA loci on the neo-Y.

**Figure 4.**
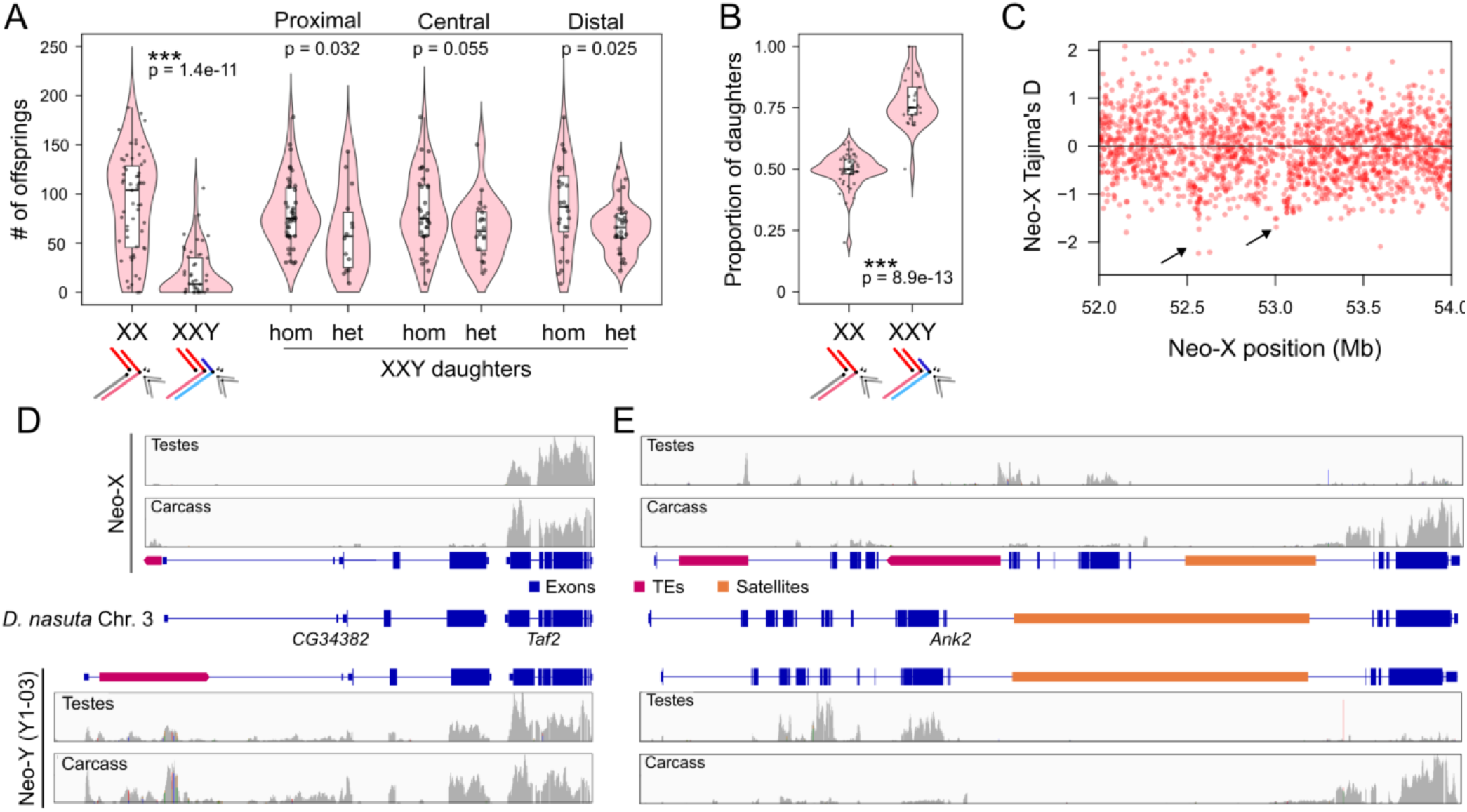
Sexual antagonistic loci and their resolution on the neo-sex chromosomes. **A**. Fecundity of euploid trivalent (XX) and aneuploid trivalent (XXY) sisters and the XXY daughters given neo-X/neo-X (hom) or neo-X/neo-Y (het) genotypes at the three molecular marker positions. P-values inferred by Wilcoxon’s rank sum test, one-tailed. **B**. Sex ratio in the offspring of XX and XXY sisters. **C**. Tajima’s D in 1KB windows at the distal end of the neo-X chromosome; arrows indicate regions with low Tajima’s D, consistent with recent selective sweeps. For values across the entire chromosome see fig. S9. **D-E**. Genome browser tracks show the genes and repeats annotations in the two regions with signatures of selective sweeps on the neo-X, their gametologs on the neo-Y, and orthologs in *D. nasuta*. Expression tracks show expression of the alleles in the testes and gonadectomized male carcass.

The female deleterious alleles on the neo-Y could either have been ancestral or accumulated due to relaxed selection after male-exclusive transmission. We reasoned that once an autosomal SA locus becomes X and Y-linked, the gametologs should experience different selective pressures; specifically, the X-gametolog will be under positive selection to lose its male-beneficial, female detrimental function, i.e. demasculination (*38, 39*). We therefore looked for signatures of selection on the distal end of the neo-X, and found two regions inside the QTL peak with markedly low Tajima’s D, consistent with recent selective sweeps (Fig. 4C). The first is *CG34382* at 52.6Mb which has high male expression but no known function; the neo-X allele acquired a DNA transposon adjacent to the transcription start site leading to the loss of expression in both the testes and male body (Fig. 4D). The second locus is at *Ankyrin2* at 53.0Mb (Fig. 4E), an isoform-rich gene known primarily for neuronal synapse stability (*40*) but also with high expression in the testes, specifically of the 5’ exons; two transposon insertions in the 5’ introns of the neo-X allele are associated with down-regulation of the testes isoform while keeping the 3’ exons unaffected. The neo-Y allele retained the ancestral state with no insertions. Both genes are consistent with positive selection for demasculinization of the neo-X gametolog. Thus, the long-range LD not only maintained the coupling of male-determining locus (the old Y) with the MBL but also prevented recombining female detrimental locus or loci onto the neo-X.

### Maintenance of long range LD favors achiasmy

The fusions creating the neo-sex chromosomes are thought to have occurred in quick succession with the neo-X arising first (*23, 25*). Given our results here, the neo-sex chromosome pair likely arose from an autosome that harbored SA loci at the distal end. Female beneficial loss-of-function allele of the SA loci then emerged and found linkage to the X via the neo-X fusion. The neo-Y fusion then followed to restore bivalent segregation which is less prone to segregation errors. Although details remain unclear on whether the SA and MBL at the distal end are the same locus and how the lineage enabled male recombination, once the SA and MBL found linkage to the Y, their great distance to the male determining locus required 2COs to maintain linkage. However, even when the chromosome is sufficiently long such that 2COs becomes more probable than 1 COs, deleterious 3 COs also become increasingly likely. As increasing genetic distance is always accompanied by decreasing non-recombinants (*41, 42*), the production of deleterious recombinants would have been both frequent and inevitable. Subsequently, shutdown of exchange between the neo-sex chromosomes emerged to simultaneously maintain long-range LD and eliminate the production of deleterious recombinant sons and daughters.

Although questions remain regarding the precise order and the molecular bases, the series of events on the neo-Y inspire unique and generalizable insights into the circumstances that predicate global recombination shutdown versus other means of suppression of exchange. The fact that local suppression via small inversions or heterochromatin formation cannot maintain long range linkage is self-evident. While large inversions can capture distal genes, effectiveness diminishes the further apart the sex-specific loci are. First, viable double COs within inversion become increasingly likely as inversion size increases, likely reducing the effectiveness of suppression of recombination (*43, 44*). Second, and more importantly, when an inversion exceeds the size of the uninverted region, it will be able to pair colinearly with the homolog (Fig. 5); at the extreme, a full chromosomal inversion is no inversion at all. Single CO within the inversion will then decouple the sex-specific allele while generating deleterious segmental duplications and deletions that affect sons and daughters, respectively, even when there is no SA. Therefore, a large inversion on the Y can be less fit than an uninverted Y even when both are equally inept at maintaining linkage between distal loci. More complex rearrangements due to multiple inversions could be suitable but are unlikely to emerge given the requirement for repeated fixations of individual small inversions with no selective mechanism to maintain their Y-linkage.

**Figure 5.**
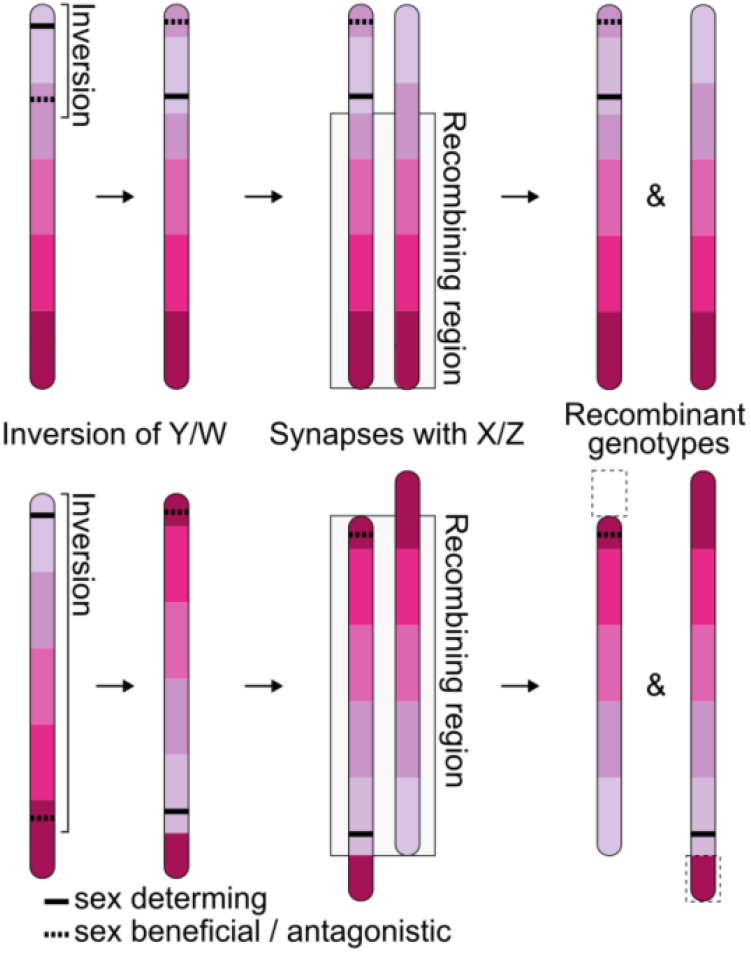
Inversion is ineffective at maintaining long range linkage. Top, when the sex-specific alleles are close, a single inversion can capture both creating an non-recombining stratum that maintains their linkage. Bottom, when the sex-specific alleles are far, a single inversion is incapable at maintaining their linkage, since synapsis can align the inverted and uninverted regions by rotating the chromosomes, allowing crossovers within the inversion, which will not only decouples the sex-specific alleles creating offspring with genotypes mismatched to their sex, but the offspring will also have segmental duplications or deletions (dotted boxes) with further deleterious effects. Note, gradient colours on chromosomes illustrate segment order.

Linkage of distal sex-specific loci thus presents a unique challenge that inversions are ill-equipped to solve. Recombination shutdown then becomes a feasible alternative. Ideally, suppression would be restricted to the sex chromosomes, limiting deleterious effects of non-disjunction in cases where COs are essential for segregation fidelity. But given minimal differentiation between the two in the incipient stages of sex chromosome formation, the homomorphic pair would look indistinguishable from autosomal pairs, making targeted suppression difficult to establish. Genome-wide shutdown, or achiasmy, is then favored, and, as previously suggested (*17*), autosomal suppression is the inadvertent pleiotropic collateral. Specifically, if the fitness advantage of maintaining linkage of male-specific loci outweighs the deleterious effects of increased nondisjunction, achiasmy can then be expected to fix. When genetic distance is long, the chance of siring unfit recombinant offspring will approach independent assortment i.e. 50%. In *Drosophila*, this is over one order of magnitude higher than the fitness cost of elevated nondisjunction rate which is typically 2-3% for achiasmate disjunction (*45*). Likelihood of achiasmy to spread will therefore depend on how stringent the requirement of COs is on segregation fidelity. Once achiasmy fixes, emergence of mechanisms to ensure faithful meiosis can be expected to follow.

## Conclusion

Prevailing models of Y chromosome evolution typically envision proximity between the male-determining and male-beneficial loci, allowing capture by inversions to establish local suppression of recombination. We presented here an unprecedented case of tight linkage between two distant male-specific loci maintained despite repeated recombination in the intervening region. This unusual linkage pattern on the young Y entails a novel scenario for recombination suppression, shedding light on the puzzling transition to achiasmy seen in the heterogametic sex.

## Notes

### Competing Interest Statement

The authors have declared no competing interest.

### Funding

- NSERC: RGPIN-2023-05390
- Michael Smith Health Research BC: SCH-2024-03818

