## Supplemental material for "Long-range linkage maintained male-specific loci on a young Y chromosome prior to recombination shutdown"

### MATERIALS AND METHODS

#### Fly stocks and strains

| Species | Strain No. | Source | Notes |
|---|---|---|---|
| *D. albomicans* | 15112-1751.03 (ref) (*1*) | NDSSC | Abbreviated as 03 |
| *D. albomicans* | kawa-012 | Kyorin/Ehime Stock Center | |
| *D. albomicans* | KBCC001 | Kyorin/Ehime Stock Center | |
| *D. albomicans* | Shl-2 | Zhang et al 2015 (*2*) | |
| *D. albomicans* | E-10857 | Kyorin/Ehime Stock Center | |
| *D. albomicans* | E-10852 | Kyorin/Ehime Stock Center | |
| *D. albomicans* | E-10809 | Kyorin/Ehime Stock Center | |
| *D. albomicans* | KM01-2 | Kyorin/Ehime Stock Center | |
| *D. albomicans* | 15112-1751.02 | NDSSC | |
| *D. albomicans* | FKC20 | Kyorin/Ehime Stock Center | |
| *D. albomicans* | OKNY2 | Kyorin/Ehime Stock Center | |
| *D. albomicans* | NG3 | Kyorin/Ehime Stock Center | |
| *D. albomicans* | KM070 | Zhou et al 2012 (*3*) | |
| *D. albomicans* | KM126 | Zhou et al 2012 | |
| *D. albomicans* | KM134 | Zhou et al 2012 | |
| *D. albomicans* | KM165 | Zhou et al 2012 | |
| *D. albomicans* | KM55 | Zhou et al 2012 | |
| *D. nasuta* | 15112-1781.00 (ref) | NDSSC | Abbreviated as 00 |
| *D. nasuta* | VNS23 | Satomura & Tamura 2016 (*4*) | |
| *D. nasuta* | 15112-1781.13 | NDSSC | |
| *D. nasuta* | 15112-1781.02 | NDSSC | |
| *D. kepuluauna* | 15112-1761.03 (ref) | NDSSC | |
| *D. sulfurigaster* | 15112-1831.01 (ref) | NDSSC | |
| *D. sulfurigaster* | 15112-1831.02 | NDSSC | |
| *D. sulfurigaster* | 15112-1811.04 | NDSSC | |
| *D. pallidifrons* | E-19901 | Kyorin/Ehime Stock Center | |
| *D. niveifrons* | LAE-276 | Kyorin/Ehime Stock Center | |

Stocks and crosses are reared on standard cornmeal media in 25 degrees 75% humidity incubators. Crosses were conducted by mating 5-8 flies of each sex.

**High molecular weight DNA extraction and Oxford Nanopore long read sequencing**
To generate assemblies of the neo-Y chromosomes which needs to be generated as a diploid phased assembly, *D. albomicans* males from different inbred lab strains were crossed with *D. nasuta* females (reference strain (*5*)) in order to increase the divergence between the neo-Y and its homolog (Chr. 3). 100-150 F1 hybrid males were collected and flash frozen. DNA extraction started following the Animal Tissue protocol for the Promega Wizard Genomic DNA Purification Kit (Catalog #: A1125). Flies were homogenized in 600ul of nuclei lysis buffer with sterile plastic pestles in batches of ~50 followed by addition of 10ul of proteinase K and incubation at 55°C for 3 hours with gentle mixing. At the isopropanol precipitation step, lysates from batches were combined in 15mL conical tubes with equal parts isopropanol. The lysate and isopropanol are gently mixed by stirring with a 20ul pipet tip and longstrand DNA fibers were spooled using the pipet tip and transferred to fresh tube for ethanol cleanup and resuspension following protocol. 03 and KBCC001 were further purified and size selected using the BluePippin using the High Pass Plus Cassette to select for 15kb+ DNA (Cata;pg # BPLUS10). For Shl-2 and kawa-012, the ONT Short Fragment Eliminator Kit (Catalog # EXP-SFE001) was used. For library preparations, ONT Ligation Sequencing Kit (SQK-LSK114) was used following the manufacturer protocol with starting material of 1.5mg of size-selected DNA. Libraries were loaded onto MinIon Flow Cells. Shl-2 was generated on the R9 chemistry while the remaining samples were generated on the R10 flow cells. Runs were terminated once 15Gb of data were generated or when the available pore counts dropped below 100. High-accuracy base calling was used.

**Phased assembly of the neo-Y chromosomes and scaffolding**
Reads were first error-corrected using the HERRO module in dorado (v1.0.2) to generate corrected reads (https://github.com/nanoporetech/dorado). Then hifiasm (0.24.0-r702) was used for phased haplotype-aware assembly (*6*). The corrected reads were used as the input and uncorrected reads were used as the ultra-long reads (--ul) with a cutoff (--ul-cut) of 10000. Illumina sequencing of the *D. nasuta* (reference strain) and males from the corresponding *D. albomicans* strains were used as the maternal (-1) and paternal (-2) genomes, respectively, for phasing. Based on Mummer (v3.23) (*7*) whole genome alignments to the published *D. albomicans* female reference (of strain 03) (*1*), we identified neo-Y contigs that align to the neo-X and evaluated the contiguity and orientation and the presence of large scale structural variations. Note, for kawa-012, the neo-Y was already assembled in its entirety at this stage and no further scaffolding was needed, but contiguity varied for the others (Table S1). We then isolated the neo-Y contigs. For Shl-2 which is the least contiguous, we additionally used samba in MaSuRCA-4.1.0 (*8*) to extend and improve the contiguity with the corrected long reads. Upon whole genome alignment with Mummer and inspection with dotplot (Fig. S4), we identified zero contigs that showed alignment characteristics diagnostic of megabase-scale rearrangements despite the high contiguity, although we cannot rule out the possibility that the contig breaks occurred precisely at inversion breakpoints. We then scaffolded the contigs based on contig overlaps when available or synteny with the neo-X and added 20 Ns between scaffolded contigs. For error correction, we performed two rounds of polishing with Hapo-G (v1.3.8) (*9*) using Illumina whole-genome sequencing reads that were trimmed with fastp (v0.22.0) (*10*). For strain 03, Illumina libraries were generated from *D.albomicans* males. For strains KBCC001, kawa-012, and Shl-2, we generated *D.albomicans* x *D.nasuta* hybrids, with *D.albomicans* as the father, for illumina DNA sequencing to reduce ambiguous read mapping due to the lack of of stain-specific neo-X assemblies.

**Gene and repeat annotation of the neo-Y assemblies**

To generate gene annotations for the assembled neo-Y chromosomes, we used LiftOn (v1.0.8) (*11*) to identify homologous neo-Y alleles/gametologs based on the NCBI genome (GCF_009650485.2) annotation which has the neo-X but not neo-Y. To classify frameshifts, we extracted the protein and coding sequences (CDSs) of genes from neo-X and strain-specific neo-Ys using gffread (v0.12.7) (*12*). Neo-Y CDS and protein sequences of each strain were aligned pair-wise with the corresponding neo-X sequences from *D.albomicans* 03 using MAFFT (v7.505) and Needle from EMBOSS (v6.6.0.0) software package, respectively. A gene was classified as having a frameshift when one or more indels changed the relative reading frame and the original frame was not restored before a stop codon was reached. To avoid indel that are artefacts, we excluded frameshift that overlap with homopolymer tracks, which are prone to sequencing errors.To annotate repeats, we used RepeatMasker (v4.1.9) with the species-group-specific repeat index from (*5*).

**RNA extraction, sequencing, and processing**

For each replicate, two 5-8 days old adults were dissected for the gonads (ovaries and testes), and the remaining carcass were collected for carcass samples. For D. albomicans male RNA-seq samples (Carcass and testes), males from the strain of interest were crossed with females from the reference strain (15112-1751.03) to reduce complications in downstream allele-specific inference. Samples were placed in Trizol, flash frozen, and stored in Trizol. Tissue was homogenized in Trizol using a motorized pestle, followed by standard RNA extraction protocol. Illumina TruSeq Stranded mRNA Library Prep kit (20020594) was used to generate RNA libraries which were subsequently sent to Novogene for QC and sequencing on the Novaseq+ 10B platform. Reads with QC'ed with fastQC (v0.12.1) and trimmed with fastp (v0.22.0), followed by mapping to sex-specific reference genome using hisat2 (v2.2.1), i.e. the neo-Y assembly was added to the reference (female) genome, for mapping of male samples. To differentiate neo-X and neo-Y mapping reads, reads were filtered using samtools (v1.17) to remove reads that map to the two equally well. Featurecount (v2.0.6) (*13*) was used for read counting at genes, using paired end (-p) and strand specific options (-s 2). Read mapping was visualized using IGV (v2.19.2) (*14*). Significant differential expression was determined using DESeq2 (v1.38.3) (*15*).

**DNA extraction and sequencing, and processing**

For each *D. albomicans* strain, 5-10 sexed flies were collected and homogenized as per the Animal Tissue protocol for the Promega Wizard Genomic DNA Purification Kit (Catalog #: A1125) for DNA extraction. Illumina TruSeq DNA Nano Kit (Catalog #: 20015964) was used for library prep with starting DNA concentration of 100ng and size fragmentation of 550bp. For single fly sequencing, Illumina DNA Prep Kit was used with up to 50ng of DNA (Catalog #: 20091660). Libraries were sent to Novogene for QC and sequencing on the Novaseq+ 10B platform. Reads were QC'ed with fastQC and trimmed with fastp (v0.22.0). For neo-Y genotyping and phylogenetic inference, all samples were mapped the the D. albomicans female reference using bwa mem (v0.7.17-r1188) (*16*), and GATK (v4.4.0.0) (*17*) haplotypeCaller was used following GATK best practices for to generate the genotype vcf file. Polymorphisms data were filtered for SNPs and bi-allelic sites using bcftools (v1.17) (*18*).

**Chromosome-scale neo-Y SNP-genotyping and phylogeny construction and inference**

As per Wei & Bachtrog 2019, neo-Y SNPs were determined by identifying sites where the female genotype is homozygous (e.g. A/A) , but the male genotype is heterozygous (e.g. A/G). The neo-Y genotype is then the polymorphic nucleotide in the male that is absent in the female (e.g. G). We then generated "pseudogenomes" of the neo-X and neo-Y of different *D. albomicans* strains, and other species by replacing the positions with SNPs based on the genotyping results. Sequences from each strain/chromosome were then extracted in non-overlapping sliding windows of 200kb and merged to generate phylogenetic trees across Muller CD using IQ-TREE (v1.6.12) with the GTR substitution model and ultrafast bootstrapping (-bb 1000). Trees were rooted by D. niveifrons, and visualized using FigTree (v1.4.4) (https://tree.bio.ed.ac.uk/software/figtree/). Tree topology inferences were conducted using the R package Phytools (v1.5.1) (*19*). Specifically, the is.monophyletic(), findMRCA(), and getSisters() functions were used to determine the extent of exchanges and relatedness between the neo-X and neo-Y genotypes and groups.

**Population genetic inference**
For genotype and haplotype representation plots (Figure 1B, and Supp figure), rare alleles were removed by minor allele frequency of 0.25; 10 random positions were selected every 50kb, and nonreference positions are plotted. We used vcftools (*20*) for sliding window nucleotide diversity and Tajima's D measurements. We first subsetted the genotype data to the appropriate sample sets, followed by the –window-pi 10000 and –TajimaD 10000 options. For chromosome-wide LD inference, we applied a minor allele frequency filter of 0.10, removing all singletons and rare alleles on the neo-Y and then subsetted the genotypes by randomly selecting 100 SNPs for every 50kb. Pair-wise LD was calculated using PLINK (v1.90b7.2) (*21*) with the settings --allow-extra-chr --r2 --ld-window-kb 10000000 --ld-window 999999 --ld-window-r2 0. For visualization, LD estimates were averaged across the window and plotted as heatmap in R.

**Identification of indel polymorphisms for molecular markers between neo-X and neo-Y**
Given the neo-Y and female assembly of strain 03, we inferred indel sites between the neo-X and neo-Y using show-snps function in Mummer (4.0.1). We then identified indel polymorphisms > 50bp at the proximal, middle, and distal locations and extracted 1kb of sequences around them. NCBI primer blast was used to identify appropriate primers around the indels show below.
Proximal primers:
5.18 MB: CCATTACGAGTGCGGTTCCT (F) and AGGCCATTAAAGGGCGTCAT (R)
Middle primers:
19.1 MB: AACTTTTAAGCGGCTTGGCG (F) and TCTGATGCCGCTTATTTCCG (R)
19.5 MB: TCGGTCCAATGGTCACGTA (F) and GAAACAGCGCCACCTTGG (R)
Distal primers:
44.8 MB: ATGCACGGCAGAAGATCGAG (F) and CGAGCCCATTGCCTCAATA (R)
45.87 MB: GGCTTGACTGTCCATGCGAT (F) and GACGCTTAAGCACAATGCCC (R)

**Generation of *yellow* mutant with CRISPR-Cas9 in D. nasuta and subsequent introgression into *D. albomicans***
We targeted three PAM sequences in the coding region of exon1 of yellow in *D. nasuta*:
CGCCGAGCAACAGCAGAAGA AGG,
ACTTTGCGTTCCCCAGCGAG AGG,
GTTCTGTGGAATGTAATCGC CGG. (PAM sequences underlined)
We followed the recommendation from the NEB EnGen sgRNA Synthesis Kit (Catalog #: E3322V) for in vitro transcription of gRNA, which were then purified using Monarch Spin RNA Cleanup Kit (Catalog #: T2040S). All gRNAs were mixed together with EnGen® Spy Cas9 NLS (Catalog #: M0646T) to a final concentration of 500 ng/ul.

For embryonic microinjections, ~200 flies were maintained in embryo collection cages and allowed to lay eggs in 30 min windows. Because the females in the species group embed the embryos deeply into the media, instead of standard agarose plates, we spread yeast paste in the collection dish for egg laying, from which the embryos were rinsed out. Microinjections were carried out by the commercial service GenomeProLab (Quebec, Canada). Yellow G1 males were identified after injecting over 1000 embryos. Yellow siblings were crossed to generate the homozygous *yellow*$^-$ line. This is the first successful CRISPR mutant generated for the species group.
To introduce *yellow*$^-$ into *D. albomicans*, *yellow*$^-$ males were crossed with *D. albomicans* females (ref strain), to generate heterozygote daughters, which are mated individually to *D. albomicans* wildtype males. As the resulting daughters are phenotypically wildtype and have equal chance of being heterozygous mutant or homozygous wildtype, each female is mated with a *D. albomicans* wildtype male. The female producing *yellow* sons at 50% frequency are thus heterozygotes, and her daughter are again mated individually with *D. albomicans* males. This process was repeated for 8 generations, followed by sib-mating to generate the *yellow* mutant line of *D. albomicans*. The karyotype of the resulting introgressed mutation was confirmed using mitotic spreads and WGS.

**Recombinant neo-Y mapping**
XXY females were generated by conducting the complex cross scheme shown in fig. S6 with the reference strains. As euploid XX and aneuploid XXY females are indistinguishable, single pair matings were conducted, and the female genotype was determined by the presence of neo-Y linked molecular markers after a two week egg lay period. XXY genotype was inferred based on the presence of neo-Y genotype at the centromere proximal location. Sons were collected and individually mated to 2 virgin *D. albomicans* 03 females. The males are collected for DNA extraction after a 2-week egg laying period, and considered sterile if no offsprings can be observed. Every male is PCR-genotyped at the proximal, middle and distal locations. To specifically select males with matrilineal neo-Y chromosomes, XXY females are crossed to *y*$^-$ *D. albomicans* males and only the *y*$^-$ sons (patrilineal X and matrilineal Y) were used for further crossing and genotyping.

**Mitotic chromosome squashes**
Larval brain mitotic squashes procedure was adapted from (*22*). Brains from sexed (based on genital imaginal discs) 3rd instar larvae were dissected in PBS, then treated with 0.5% sodium citrate for 8 minutes. Samples were fixed in paraformaldehyde, diluted to 1.8% with nuclease-free water and glacial acetic acid, for 6 minutes before transferring onto a Sigma-coated coverslip and standard slide. Firm pressure was applied perpendicular to the slide surface to squash the larval brains. Slides were subsequently submerged in liquid nitrogen until vigorous bubbling ceased. Coverslip was immediately flicked off using a razor blade and slides were submerged in ethanol for at least 30 minutes before mounting in Slowfade Antifade Mountant (Thermofisher # 361055759). Mitotic squashes of stock strains, F1, and F2 were performed to confirm expected karyotypes and identify aneuploid offspring karytypes. Larval brain mitotic spreads were captured using the Zeiss Axio Observer 7 with Apitome 3 at 63x. Images were deconvoluted with Apitome plus processing in Zeiss's Zen software.

**Fluorescent imaging of testes**
For each slide, 3 pairs of testes were dissected from male flies in PBS and fixed in 4% paraformaldehyde diluted with PBS for 15 minutes at room temperature. Three 5 minute washes of PBST-X (0.1% Tween20, 0.2% Triton-X) were performed at room temperature and followed by two 15 minute washes of 0.3% sodium deoxycholate. Samples were incubated for 1 hour at room temperature in 3% BSA-PBST with DAPI added in a 1:2000 ratio. Samples were washed in PBST-X three more

times prior to mounting in Slowfade Antifade Mountant. Testes images were captured using the Olympus IXplore IX85 Spin. Whole testes image was taken at 25x magnification by tiling images across the whole length of the testes via the Multiple Image Alignment acquisition feature. Specific sperm bundles and testis morphologies were obtained at 60x magnification with immersion oil applied using the maximum intensity projection of Z-stacks. Images were processed using Olympus's CellSens software.

### DATA AVAILABILITY

Raw reads have been deposited on the Sequence Read Archive under PRJNA1521485.

### SUPPLEMENTARY FIGURE LEGENDS

**Figure S1.** Representative phylogenetic trees of Muller CD genotypes across the species group. Individual neo-Ys are coloured in shades of blue, and neo-Xs are coloured in red. The neo-Ys that were assembled are indicated by stars.

**Figure S2.** Genotype representation of neo-X and neo-Y of sequenced inbred lines. For each line, the upper track represents the neo-X and the lower track represents the neo-Y.

**Figure S3.** Neo-Y scaffolding and alignments to Neo-X. Neo-Y contigs aligned to the neo-X shown in dotplots. Horizontal lines delimit contig breaks and points of connection. Stars indicate regions with contig overlaps that are merged.

**Figure S4.** Neo-X and neo-Y expression difference in testes (left) and gonadectomized whole body (right). All males are crossed to reference females. Significantly differentially expressed genes are shown in blue. Genes exceeding log2(fold-difference) of ±10 are shown in triangles.

**Figure S5.** Cross scheme to generate neo-Y recombinants. A. Multi-generational crosses generating XXY females. B. We generated a mutant line of the X-linked gene *yellow* with CRISPR-Cas9 – the first time in the species group – that allows differentiation between patrilineal and matrilineal transmission of the Y chromosome. *yellow* mutant and wildtype males are shown. C. Punnett square showing all possible offspring genotypes from XXY females. Lethal genotypes and sex are indicated.

**Figure S6.** Haplotype inference of whole genome sequenced recombinant males. Genotype inference distinguishes neo-X (pink), neo-Y (blue), and Chr. 3 (gray) genotypes. First individual is a recombinant son produced by an XX trivalent, shown for illustrative purpose. Regions of the genome with only one colour represent homozygosity.

**Figure S7.** Offspring genotype frequencies from crossing XXY and XX trivalent sisters to yellow males. Expected frequencies derived from cross scheme in fig S5, assuming equal transmission of all possible gamete configurations and no lethal genotypes.

**Figure S8.** Tajima's D across the entire neo-X chromosome. Each point represents a 1kb window. Horizontal bar indicates the region shown in Figure 4C.
