## Supplemental figures for "Long-range linkage maintained male-specific loci on a young Y chromosome prior to recombination shutdown"

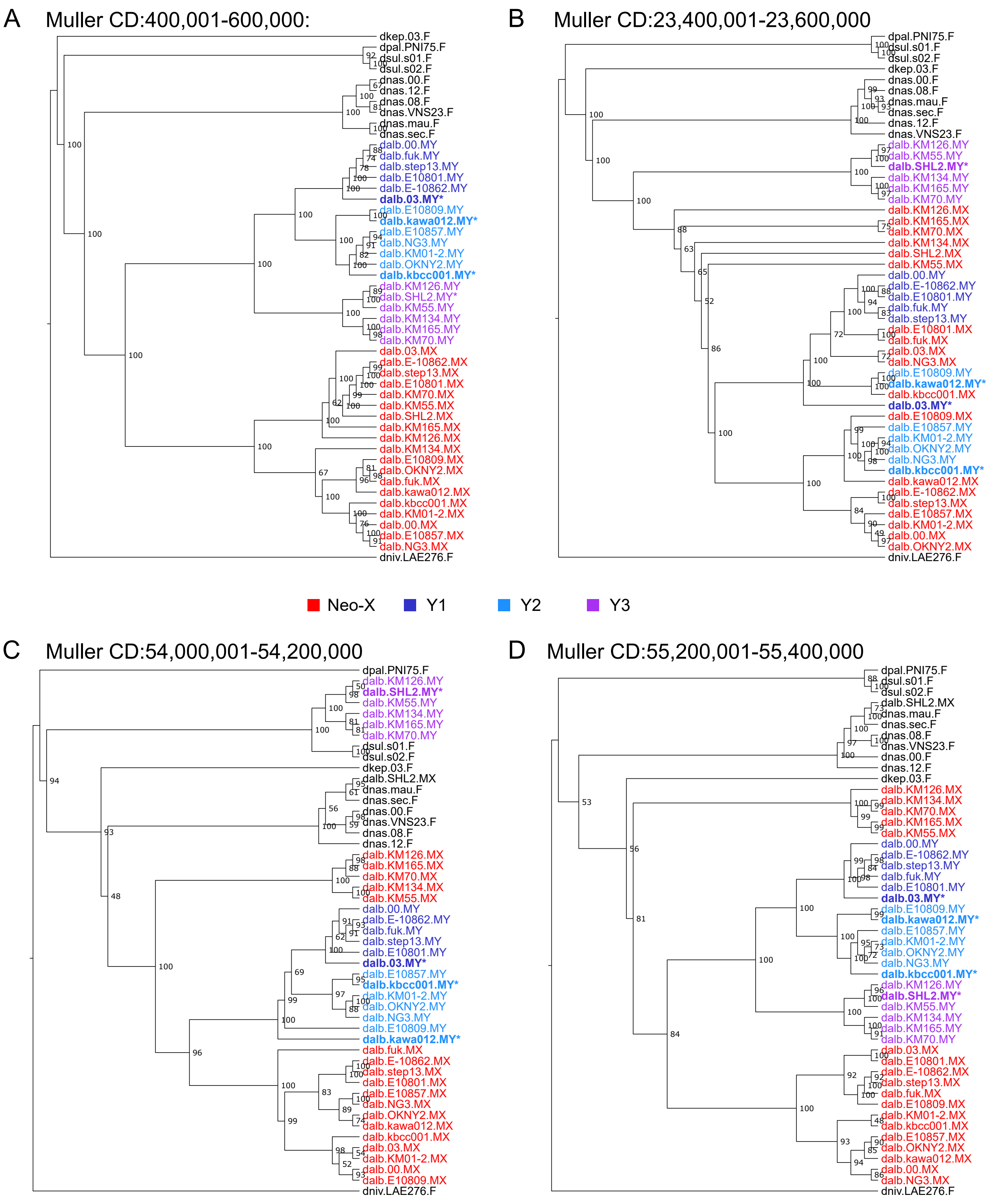


Figure S1. Representative phylogenetic trees of Muller CD genotypes across the species group. Individual neo-Ys are coloured in shades of blue, and neo-Xs are coloured in red. The neo-Ys that were assembled are indicated by stars.

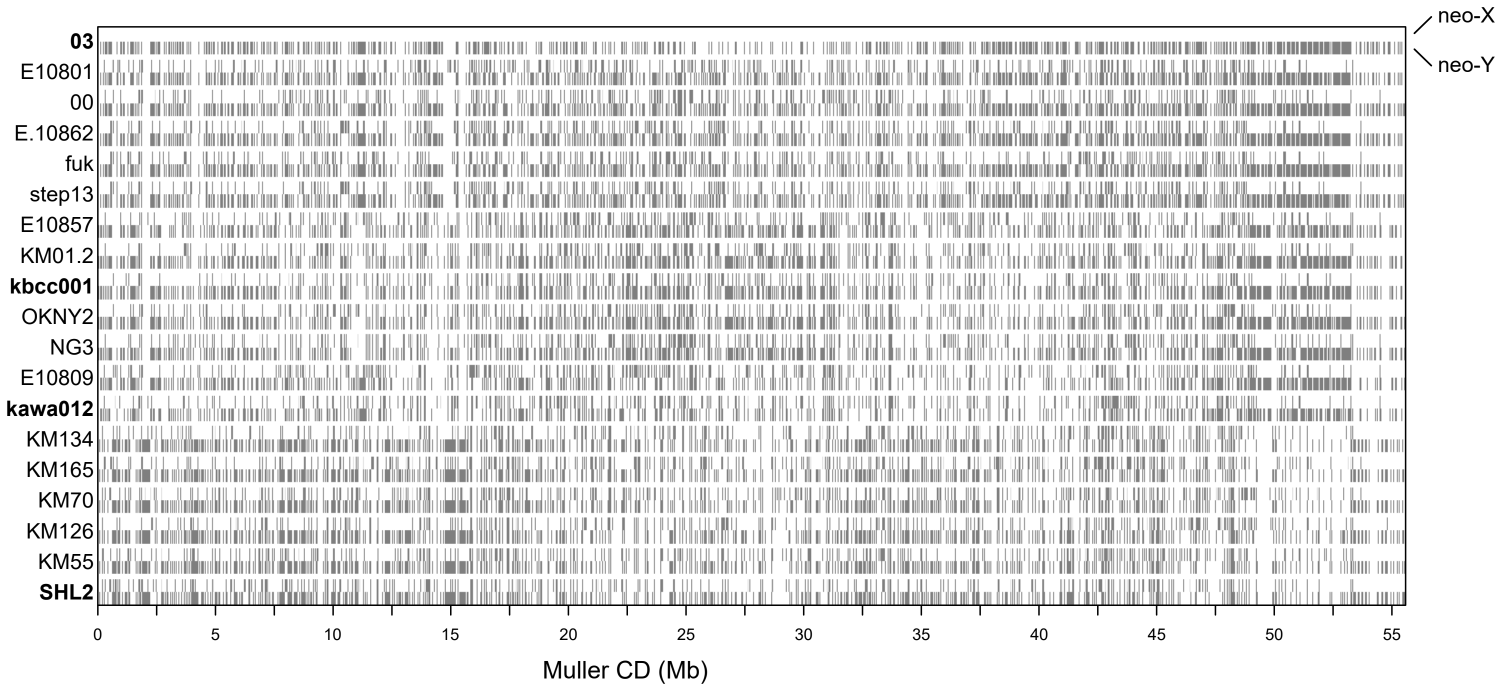


Figure S2. Genotype representation of neo-X and neo-Y of sequenced inbred lines. For each line, the upper track represents the neo-X and the lower track represents the neo-Y.

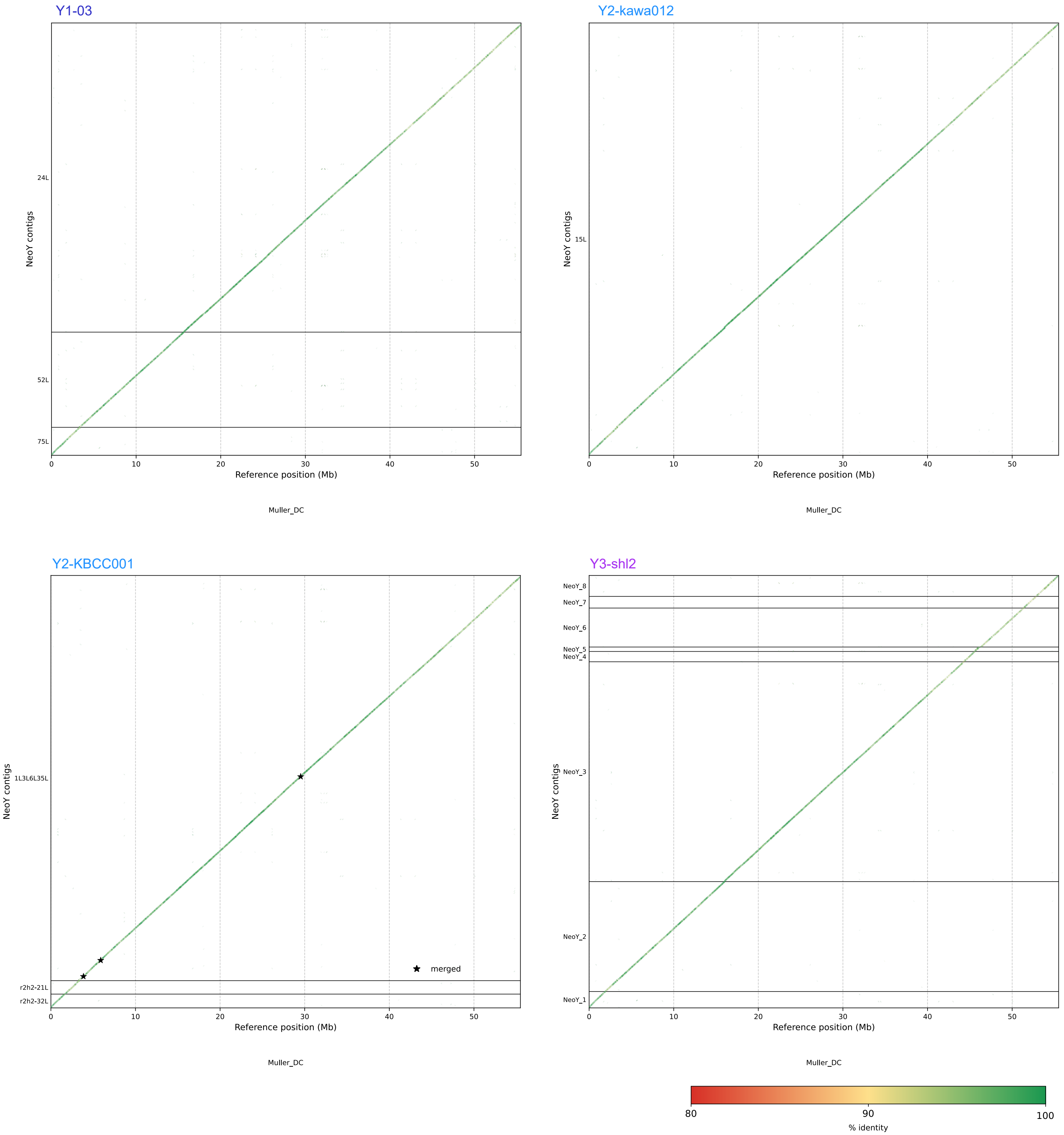


Figure S3. Neo-Y scaffolding and alignments to Neo-X. Neo-Y contigs aligned to the neo-X shown in dotplots. Horizontal lines deliminate contig breaks and points of connection. Stars indicate regions with contig overlaps that are merged.

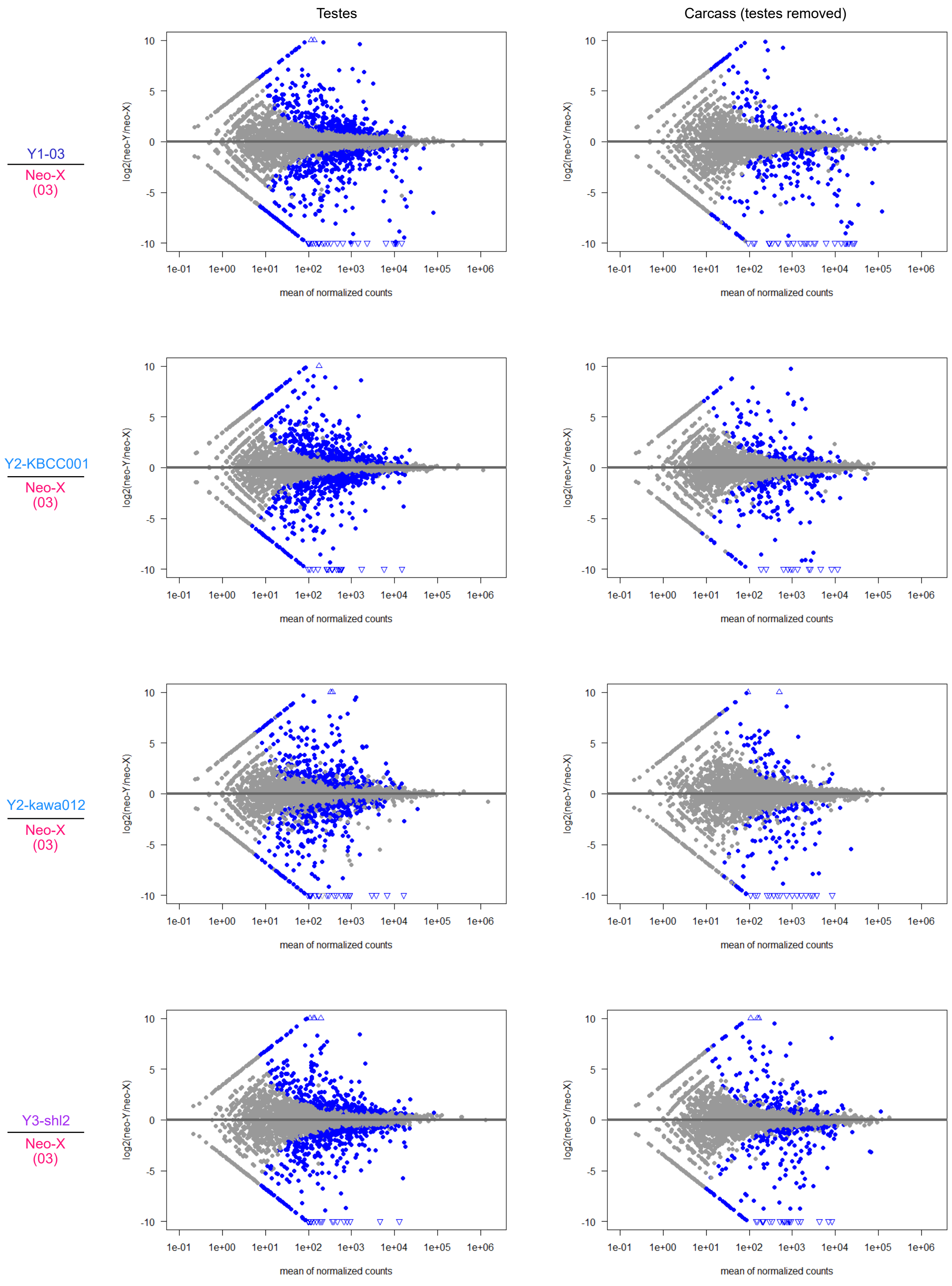


Figure S4. Neo-X and neo-Y expression difference in testes (left) and gonadectomized whole body (right). All males are crossed to reference females. Significantly differentially expressed genes are shown in blue. Genes exceeding log2(fold-difference) of ±10 are shown in triangles

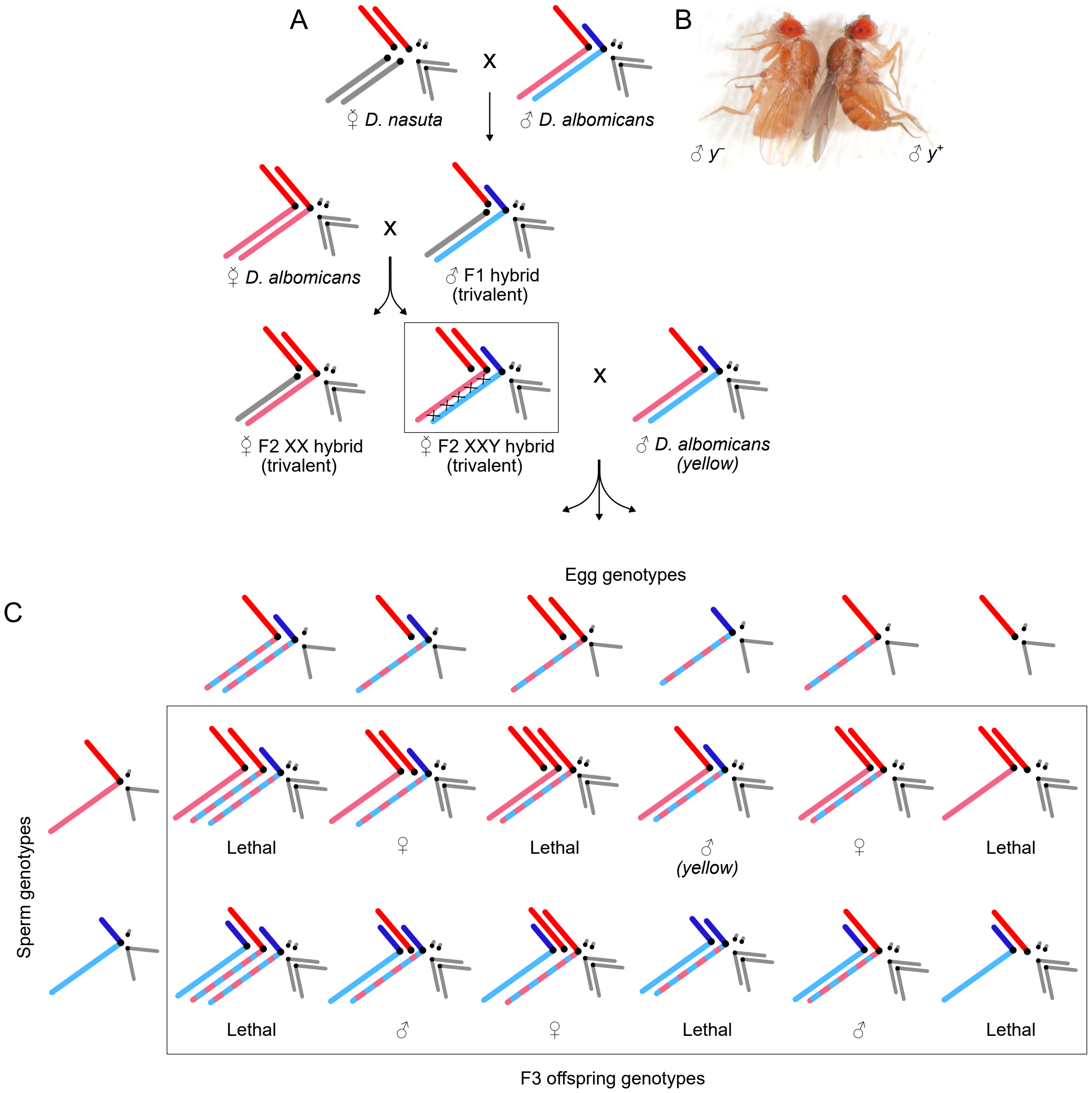


Figure S5. Cross scheme to generate neo-Y recombinants. A. Multi-generational crosses generating XXY females. B. We generated a mutant line of the X-linked gene yellow with CRISPR-Cas9 – the first time in the species group – that allows differentiation between patrilineal and matrilineal transmission of the Y chromosome. Yellow mutant and wildtype males are shown. C. Punnett square showing all possible offspring genotypes from XXY females. Lethal genotypes and sex are indicated.

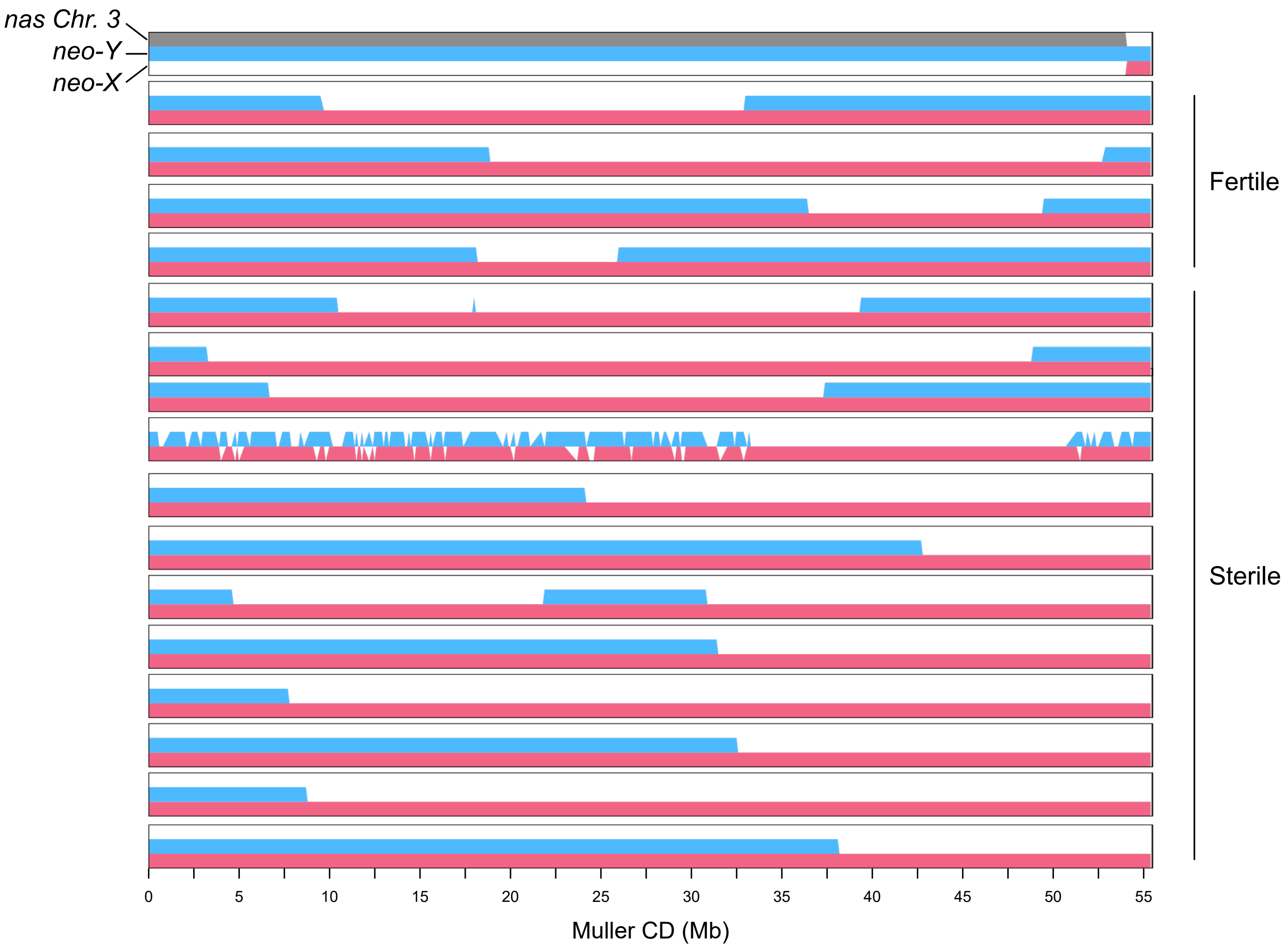


Figure S6. Haplotype inference of whole genome sequenced recombinant males. Genotype inference distinguishes neo-X (pink), neo-Y (blue), and Chr. 3 (gray) genotypes. First individual is a recombinant son produced by an XX trivalent, shown for illustrative purpose. Regions of the genome with only one colour represent homozygosity.

| | | Daughters | Sons | | Total |
|---|---|---|---|---|---|
| | | | *WT* | *yellow* | |
| 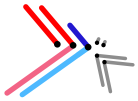 | XXY trivalent (n = 48) | 689 (75.5%) | 150 (16.4%) | 74 (8.1%) | 913 |
| | XXY trivalent expected | 50.0% | 33.3% | 16.6% | |
| 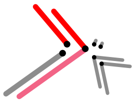 | XX trivalent (n = 76) | 2662 (50.2%) | 2618 (49.4%) | 23 (0.4%) | 5303 |
| | XX trivalent expected | 50.0% | 50.0% | 0.0% | |

Figure S7. Offspring genotype frequencies from crossing XXY and XX trivalent sisters to *yellow* males. Expected frequencies derived from cross scheme in fig S5, assuming equal transmission of all possible gamete configurations and no lethal genotypes.

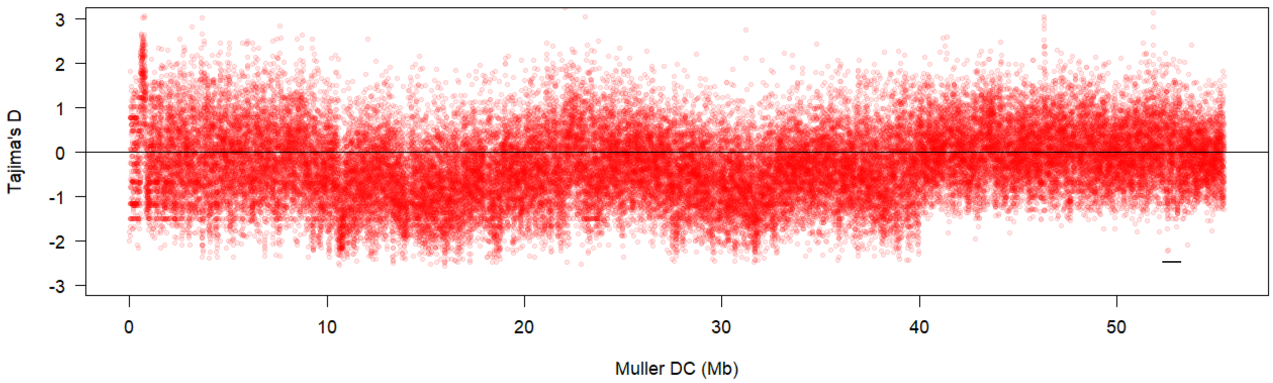


Figure S8. Tajima's D across the entire neo-X chromosome. Each point represents a 1kb window. Horizontal bar indicates region the shown in Figure 4C.
